# Intact double stranded RNA is mobile and triggers RNAi against viral and fungal plant pathogens

**DOI:** 10.1101/2021.11.24.469772

**Authors:** Christopher A. Brosnan, Anne Sawyer, Felipe F. de Felippes, Bernard J. Carroll, Peter M. Waterhouse, Neena Mitter

## Abstract

Topical application of double-stranded RNA (dsRNA) as RNA interference(RNAi) based biopesticides represents a sustainable alternative to traditional transgenic, breeding-based or chemical crop protection strategies. A key feature of RNAi is its ability to act non-cell autonomously, a process that plays a critical role in plant protection. However, the uptake of dsRNA upon topical application, and its ability to move and act non-cell autonomously remains debated and largely unexplored. Here we show that when applied to a leaf, unprocessed full-length dsRNA enters the vasculature and rapidly moves to multiple distal below ground, vegetative and reproductive tissue types in several model plant and crop hosts. Intact unprocessed dsRNA was detected in the apoplast of leaves, roots and flowers after leaf application and maintained in subsequent new growth. Furthermore, we show mobile dsRNA is functional against root infecting fungal and foliar viral pathogens. Our demonstration of the uptake and maintained movement of intact and functional dsRNA stands to add significant benefit to the emerging field of RNAi-based plant protection.

---

RNA interference (RNAi) represents one of the frontline defences plants utilise in combating a wide range of pests and pathogens. At the centre of RNAi is the processing of long double-stranded RNA (dsRNA) into 21-24 nucleotide (nt) small interfering RNA (siRNA) by DICER-like (DCL) enzymes. These siRNAs are subsequently loaded into ARONAUTE (AGO) proteins and via their complementarity with target RNA or DNA, illicit their silencing function. The plant biotechnology field has harnessed this natural defence mechanism to protect crops from a variety of plant pests and pathogens using genetic modification (GM) (1, 2). More recently, the adaptation of RNAi in the form of topically applied RNAi-based biopesticides has gained increasing traction as an effective and sustainable non-GM method of plant protection. One of the unique features of plant and some metazoan RNAi is its ability to act non-cell autonomously, moving and functioning both over short (cell-to-cell) and long distances(3). Application of dsRNA to plant tissues including leaves, roots, flowers and fruit, has been shown to initiate efficient RNAi-based protection against many plant pathogens and pests in the same localised tissue types (4–9). While this approach has demonstrated enormous potential for sustainable crop protection, the capacity of this topical form of RNAi to move cell-to-cell or over longer distances like its endogenous and transgenic counterparts remains largely unexplored. Although several reports have eluded to the potential of dsRNA to act non-cell autonomously (9–11), a clear demonstration and in-depth mechanistic insights into the process in intact plants is still lacking. Addressing this knowledge gap is key to overcoming multiple current issues in the field, such that broad spectrum, multi-tissue and sustained plant protection can be achieved.

Here we demonstrate that once applied to a plant, dsRNA moves as an intact, and subsequently maintained, unprocessed molecule to multiple distal tissues in a variety of model plant and crop species. This mobility delivers dsRNA to the shoot apex, roots and reproductive tissues, all key targets when dealing with plant pathogens. We show that movement occurs via the apoplastic channel in contrast to canonical RNAi movement which transverses cell types via the symplasm. Finally and crucially, we demonstrate that mobile dsRNA is functional against fungal and viral pathogens in several distal tissue types. Together these novel findings on the mobility and action of topically applied RNAi open wide a range of future prospects for the field as a whole.

To investigate if topically applied dsRNA could move from the site of application to distal tissue types, we chose to start in the model plant *Arabidopsis thaliana*. We assayed the potential transfer of dsRNA from its application site (i.e., the upper leaf epidermis) to the vasculature (Fig. 1a middle panel). To avoid artefacts associated with fluorophore conjugated RNA approaches, we utilized meselect (12). This technique physically separates epidermal and vascular cells and has previously been used to diagnose siRNA movement (13, 14). Using northern blot analysis, we were readily able to detect full-length dsRNA in the vasculature of the applied leaf (Fig. 1a middle panels), an essential precursor to any potential movement to distal tissue types. Next, we looked at distal tissues, including the shoot apex and roots. As early as one hour post application we could detect full-length dsRNA in both the shoot apex and the root (Fig. 1a lower panels). We then focused on other above ground and reproductive tissues of Arabidopsis, namely the stems, flowers, and siliques. Again, we were able to detect full-length dsRNA in the stems and the flowers 24 hours post application (Fig. 1a top panels). Interestingly, we not only saw mobile dsRNA within siliques already formed at the time of application, but also four days post application in newly formed siliques which were flowers at the time of dsRNA application (Fig. 1a top panels). As opposed to reproductive or root tissues, movement to pre-established young leaves was barely detectable (Fig. S1).

**Figure 1.**
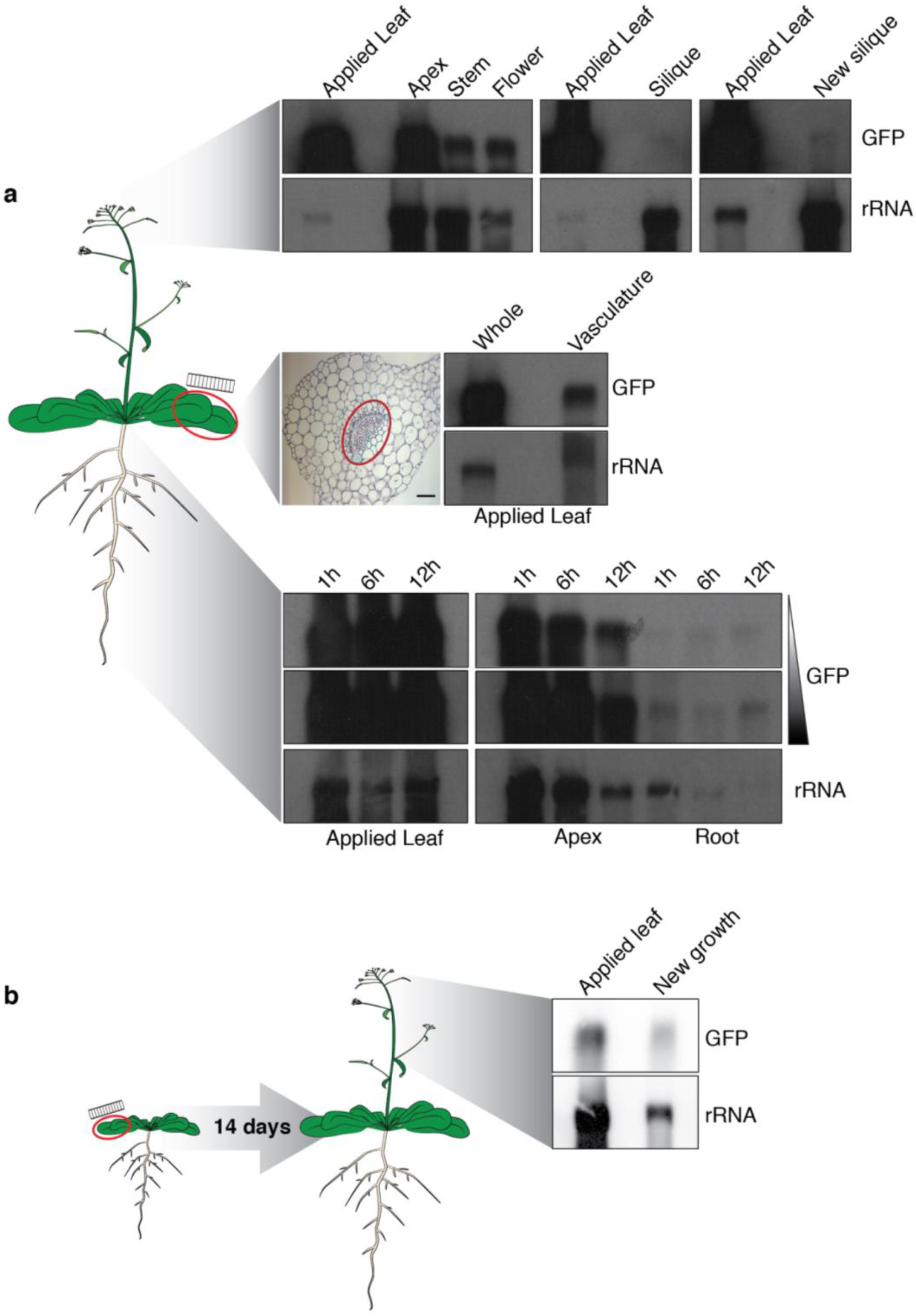
Movement and maintenance of intact topically applied dsRNA to multiple tissue types of Arabidopsis. **a,** Schematic of Arabidopsis plant and detection of mobile dsRNA in distal tissue types. Middle panels – cross section of leaf showing vasculature and RNA gel blot probing *GFP* dsRNA on the applied leaf (whole) and in the physically separated vascular tissue (vasculature). Two exposures are shown. Top panels - RNA gel blot probing dsRNA in the apex, stem, flower and silique tissues 24 h after application. Lower panels - RNA blot probing dsRNA in the apex and roots 1 h, 6 h and 12 h after application. Scale bar represents 100 μm **b**, Schematic showing growth stage of the plant when dsRNA was applied and 14 days later when dsRNA was detected in new aerial growth. Right panel - RNA gel blot probing dsRNA in new growth. Ribosomal RNA (rRNA) serves as a loading control.

The presence of dsRNA within siliques formed post-RNA application (Fig. 1a) could conceivably be due to either (a) movement into these tissues post-formation, (b) maintenance of a mobile pool of dsRNA during the transition from flower to silique, or (c) a combination of both. To test the idea that dsRNA could potentially be maintained within newly formed tissue, we extended the timeframe between application and detection to 14 days (Fig. 1b). We were able to detect dsRNA in newly formed stems and inflorescences (Fig. 1b), likely indicating that the pool of dsRNA present from previous or continued movement (e.g., to the shoot apex) was maintained during the formation of new tissue types (hereafter we refer to this as maintenance rather than movement).

By using the same experimental design in canola (*Brassica napus*) but extending the detection time to 72 hours post-application, we investigated if the observed movement in Arabidopsis was possible in another Brassicaceae species. We readily detected mobile dsRNA in canola roots, but not within young leaves (Fig. 2a). Flowering canola plants displayed a significant amount of movement from applied leaves into both the stems and floral organs, albeit movement levels were contingent on the floral node (Fig. 2a; 1-oldest, 3-youngest). Overall, this pattern of movement was similar to that seen in Arabidopsis (Fig. 1).

**Figure 2.**
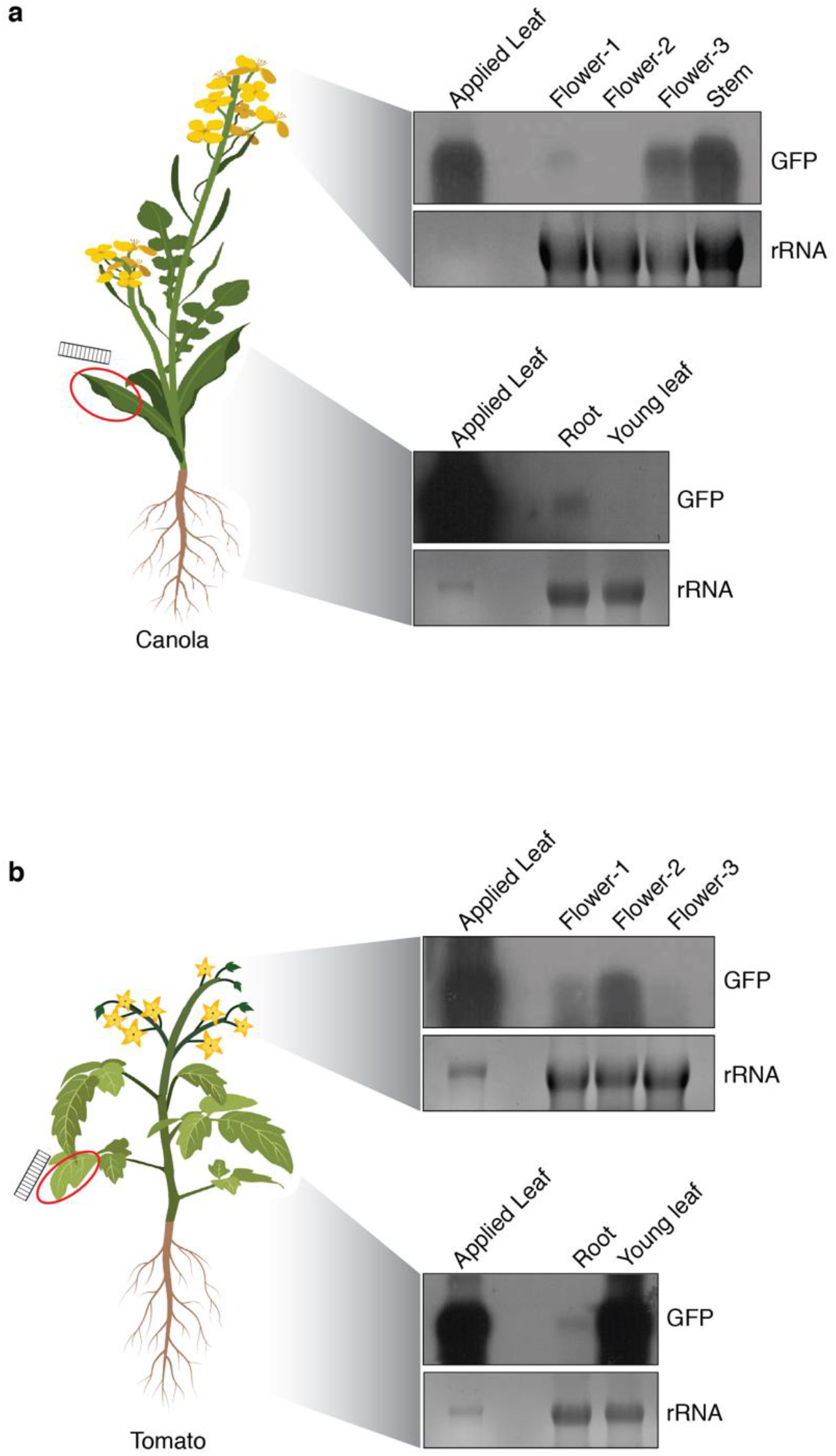
dsRNA moves to multiple tissue types in canola and tomato. Schematic of respective experimental setup is shown on the left. **a**, RNA gel blot detection of mobile dsRNA canola. Top panel - RNA blot probing *GFP* dsRNA in the applied leaf, flower-1, −2 and −3 (3 being the most apical) and stem of flowering canola plants. Bottom panel - RNA blot probing dsRNA in the applied leaf, root and young leaf of canola. **b**, RNA gel blot detection of mobile dsRNA in tomato. RNA gel blot analysis of dsRNA from the applied leaf, flower-1, −2 and −3 (3 being the most apical) of flowering tomato plants (upper panel). Lower panel - detection of RNA in applied leaf, root and young leaf of tomato. Ribosomal RNA (rRNA) serves as a loading control.

To assess whether different plant architectures influenced mobility, we shifted our focus to two model Solanaceae plant species, tomato and *Nicotiana benthamiana*. While we found moderate to low levels of detectable dsRNA in the roots of tomato plants 48-72 hours post-application, a significantly larger amount was detected in young leaves (Fig. 2b). We were also able to detect a significant amount of mobile RNA in the floral organs, again contingent on the node (Fig. 2b; 1-oldest, 3-youngest). Only barely detectable amounts of dsRNA were present in the distal young leaves of *N. benthamiana* plants (Fig. S2), which may be due to a thick waxy cuticle and/or high trichome density preventing uptake through epidermal and mesophyll cells.

Current knowledge of the channels involved in RNA movement within plants is mostly based on the movement of either transgenic or endogenous siRNAs and miRNAs (15). Both transgenic and endogenous siRNA species have long been thought to move via symplastic channels, through plasmodesmata that link the cytoplasm of adjoining cells (16–18). The most direct evidence for this channel of movement exists for the endodermis to the stele mobility of miR165 in Arabidopsis roots (19, 20). By spatially and temporally inducing deposition of callose to block plasmodesmata, the symplastic transport of miR165 could be blocked (20). We wanted to investigate whether the movement of topically applied dsRNA to distal tissue types followed a similar channel to siRNA/miRNAs (i.e. though the cytoplasm via a symplastic pathway), or if they moved more akin to nutrients and/or water (i.e. through the cell wall via an apoplastic pathway – Fig. 3a).

**Figure 3.**
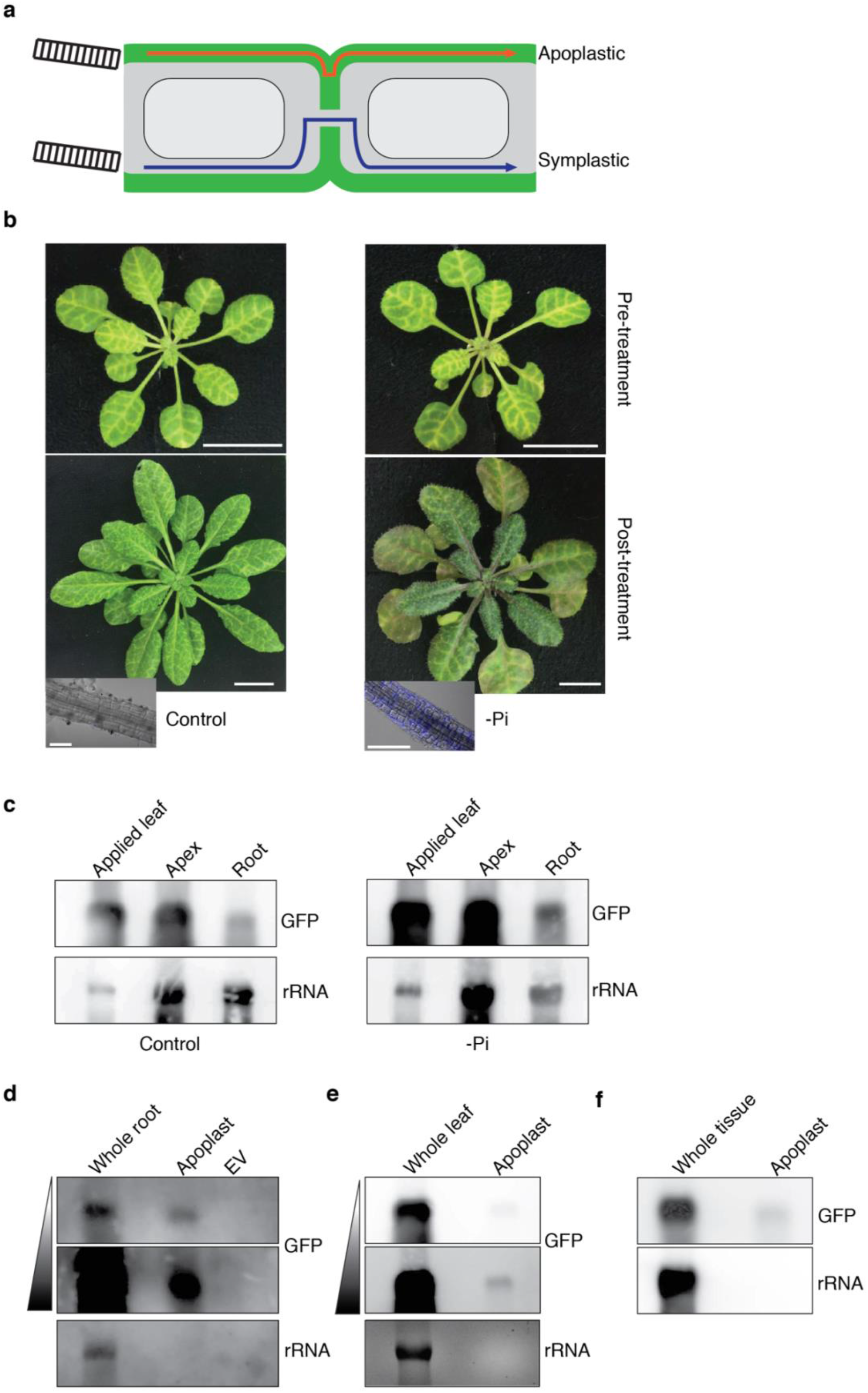
Apoplastic transport and maintenance of dsRNA. **a,** Schematic depicting two possible routes of dsRNA movement, through the cell wall via the apoplastic pathway (red line), or through the cytoplasm via plasmodesmata (symplastic pathway, blue line). **b,** Representative images of phosphate replete (left panel) and phosphate deprived (-Pi; right panel) *pSUC::SUL (SS)* plants in which callose deposition has been induced. Top panels show plants pre-treatment and lower panels show the same plants two weeks post-treatment. Lack of vein centred bleaching indicates a lack of symplastic siRNA movement. Inset image is of roots from the same plants stained for callose deposition. Scale bars represent 100 μm for roots and 1 cm for whole plants. **c**, Detection of dsRNA in the application leaves and distal apex or root tissue of control (left panel) and -Pi treated (right panel) *SS* plants. **d**, RNA gel blot analysis of apoplastic fluid and extracellular vesicle (EV) purifications from Arabidopsis distal roots containing mobile dsRNA. Two exposures are shown for the *GFP* probe. **e**, RNA gel blot analysis of apoplastic fluid purifications from distal leaves of tomato plants 48 h after RNA application to source leaves. Two exposures are shown for the *GFP* probe. **f**, RNA gel blot analysis of apoplastic fluid purifications from new growth two weeks after application of dsRNA to source leaves of Arabidopsis plants. rRNA serves as a loading control.

To address the possibility of dsRNA mobility occurring via the symplastic pathway, we blocked plasmodesmata via callose deposition. Phosphate starvation (-Pi) has previously been shown to constitutively induce callose deposition and block symplastic movement in Arabidopsis roots (21). To ensure effective blocking of plasmodesmata-based RNA transport, we conducted our experiments in the *pSUC::SUL (SS)* RNAi Arabidopsis line, which expresses a phloem companion cell-specific RNAi transgene spawning 21- and 24-nt siRNAs (22) which target the *MAGNESIUM CHELATASE (SUL)* mRNA. This line is an effective sensor of siRNA movement, with the 21-nt class diagnosing mobile RNAi as vein-proximal leaf bleaching 10-15 cells from their expression domain (22). As anticipated, we observed that phosphate replete *SS* plants displayed only background callose staining in roots and the characteristic vein-centred bleaching associated with functional siRNA movement (Fig. 3b and Fig. S3). Comparatively, two weeks after *SS* plants were deprived of phosphate (-Pi), we observed heavy callose staining in roots, with newly emerging leaves (~8 leaves) displaying a suppression of the chlorosis phenotype (Fig. 3b and Fig. S3). Together, these findings diagnose an effective blocking of symplastic siRNA movement (Fig. 3b), despite comparable levels of *SS* siRNA being present in control and -Pi plants (Fig. S3). Next, we applied dsRNA to the leaves of *SS* plants and tested if movement to the apex and roots was affected by callose-based blocking of plasmodesmata. As expected, efficient movement of dsRNA from the applied leaf to the apex and the root occurred 24 hours after application under normal growth conditions (Fig. 3c). Interestingly, we could detect equivalent amounts of mobile dsRNA (in comparison to the applied leaf) in both the apex and roots of the -Pi treatment, despite a clear lack of *SS* siRNA movement (Fig. 3b-c). This strongly indicated that, unlike siRNAs, topically applied dsRNA does not travel via the symplasm.

The other transport pathway in plants that dsRNA could conceivably travel through and be maintained in, is the apoplast. Previous work using *CYP3*-dsRNA detected fluorescence primarily within the apoplast but also in select phloem, phloem companion and trichome cells (9). Again, avoiding fluorophore-labelled RNA, we isolated apoplastic fluid from the destination tissue types of mobile dsRNA in multiple plant species to assay for dsRNA presence, we were able to detect intact mobile pools of dsRNA in the apoplastic fluid of Arabidopsis roots (Fig. 3d). While we and others (17, 18) have shown that siRNAs likely move in plants via the symplasm, it has been reported that endogenous and exogenous siRNAs are also present within extracellular vesicles (EVs; (23, 24)). Present within the apoplast, EV’s have been shown to facilitate the transfer of their siRNA/miRNA cargo to infecting fungi (24). To explore the possibility, albeit unlikely, that dsRNA is transported within the apoplast as cargo of EVs, we isolated EVs from the roots of Arabidopsis plants following application of dsRNA to the leaves. While dsRNA was detected in the apoplast, dsRNA within EVs isolated from the same apoplastic fluid, were below the detection limit (Fig. 3d). Next, we tested whether dsRNA moving to aerial tissue within an alternative species were also present in the apoplastic fluid. Using the young destination leaves of tomato, which we previously saw as an efficient sink for dsRNA movement (Fig. 2b), we were readily able to detect dsRNA in the apoplastic fluid (Fig. 3e). Finally, we tested whether the dsRNA we observed as ‘maintained’ in new growth (Fig. 1b), was also contained within the apoplast of this tissue. Extraction of apoplastic fluid from the new growth, 14 days post-application of dsRNA to source leaves, revealed a maintained pool of RNA (Fig. 3f). A clear consequence of dsRNA being translocated and stored within the apoplast would be a lack of DCL processing into small RNAs, a process requiring entry into the cell. Indeed, we were unable to detect siRNA production within tissues containing mobile dsRNA despite detecting graft-transmitted transgenic and endogenous mobile siRNA species (Fig. S4). We found that all iterations of dsRNA movement (i.e., in different species, tissue and contexts) and movement types (strict movement or maintenance) occur predominantly, if not exclusively, via the apoplastic pathway.

Having established that intact dsRNA can move to, and be maintained in multiple tissue types in a variety of plant species, we investigated whether dsRNA could effectively target pathogens. *Fusarium oxysporum* and *Verticillium dahliae* are both well characterised root-infecting fungal pathogens which cause large scale losses in a wide variety of crop species. Using transgenic isolates of each fungus constitutively expressing *GFP* (*pGDPA-GFP* or *pTEF1-GFP*), we tested if mobile apoplastic dsRNA efficiently targets fungal mRNA (Fig. 4a). Compared with control plants, we found a reproducible, ~50% reduction in *GFP* transcripts in *F. oxysporum*-infected roots upon treatment of leaves 4 dpi with dsRNA targeting *GFP* (Fig. 4b). Mobile dsRNA exhibited a similar level of effectiveness when *GFP*-expressing *V. dahliae*-infected plants were treated (Fig. 4c).

**Figure 4.**
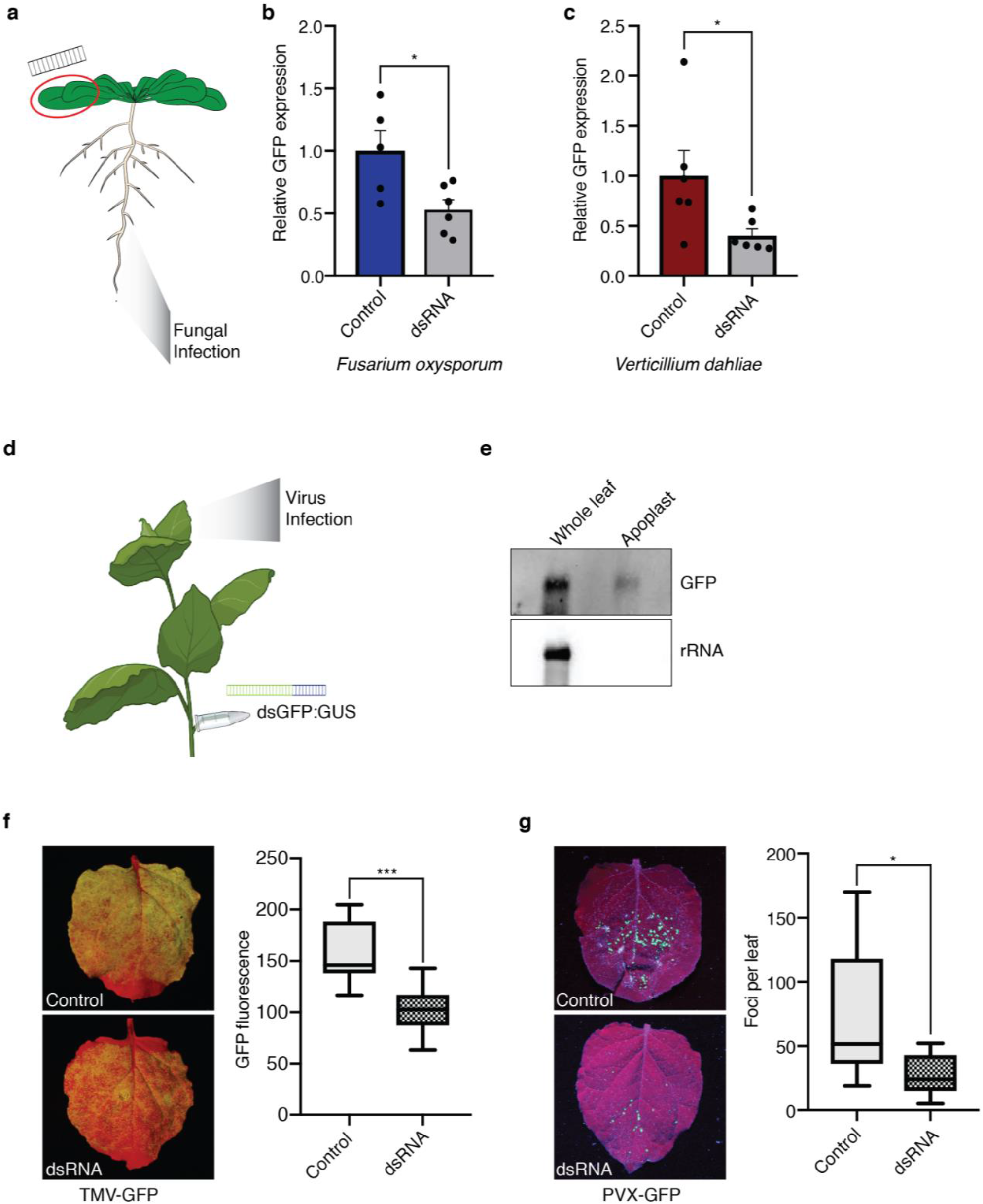
Functionality of mobile dsRNA against multiple pathogens. **a,** Schematic showing experimental setup for assaying mobile dsRNA in Arabidopsis targeting infecting fungi in the roots. **b**, Quantification of *GFP* transcript level in *pGDP-GFP* expressing *Fusarium oxysporum* from control and *GFP* dsRNA-treated Arabidopsis plants. Columns represent mean +/-s.e.m; n = 5 and 6 biological replicates respectively for each data point; Two-sided unpaired students *t*-test \**P* ≤0.05. **c**, Quantification of *GFP* transcript level from *pTEF1-GFP Verticillium dahliae* treated with control or mobile *GFP* dsRNA. Columns represent mean +/-s.e.m; n = 5 and 6 biological replicates respectively for each data point; Two-sided unpaired students *t*-test \**P* ≤0.05. **d**, Schematic of dsRNA delivery by ‘petiole absorption’ and subsequent viral infection of *N. benthamiana.* **e**, RNA gel blot detection of dsRNA in mobile leaf tissue in ‘whole leaf’ or apoplastic fluid extracted from a distal leaf of ‘petiole absorption’ *N. benthamiana* plants. **f**, Representative leaves of TMV-GFP infected *N. benthamiana* plants from either control or dsRNA petiole treated plants and quantification of GFP fluorescence of control and treated plants. Error bars represent +/- s.e.m; n = 10 leaves from 5 plants; Two-sided unpaired students *t*-test \*\*\**P* ≤0.0005. **g**, Representative leaves of PVX-GFP infected *N. benthamiana* plants from either control or dsRNA petiole treated plants and quantification of number of GFP foci of control and treated plants. Error bars represent +/- s.e.m; n = 8-10 leaves from 8-10 plants; Two-sided unpaired students *t*-test \**P* ≤0.05.

The mobility of RNAi via virus-derived siRNA (vsiRNA) has long been hypothesised to act as a defensive front against viruses, which themselves actively move throughout plants (1). To investigate if mobile topically applied dsRNA could target invading viruses, we focused on two well characterised GFP-tagged viruses, tobacco mosaic virus (TMV-GFP) and potato virus X (PVX-GFP), allowing us to monitor very early stages of infection, likely to be key in any effect seen by topical application of dsRNA (5, 25). To allow for the detection of primary siRNAs derived exclusively from exogenous dsRNA application and not secondary amplified pools of siRNA derived from the virus itself, we used a dsRNA sequence containing a portion of non-targeting RNA in addition to *GFP* (Fig. 4c; *dsGFP:GUS*). While the ideal host for each virus was *N. benthamiana*, dsRNA uptake is limited in this species (Fig. S2). Therefore we chose to use petiole uptake, which has been shown to effectively translocate dsRNA from the source petiole to the apical sink leaves (10). Key to this was that the dsRNA movement afforded by petiole absorption was reportedly via the same apoplastic channel as we had observed for topical application. This was indeed the case, with petiole-absorbed dsRNA detected within the apoplast of the young destination leaves (Fig. 4e). Infection with movement compromised TMV-GFP 3-four days post-dsRNA application resulted in a reproducible decrease in GFP-fluorescence indicating a decrease in viral load when compared to the control (Fig. 4f). We next conducted a similar experiment, this time using PVX-GFP. A significant decrease in individual infection foci was seen upon mobile dsRNA treatment, highlighting the functionality of the mobile pool of dsRNA 4 dpi/7 days post dsRNA application (Fig. 4g). Despite the clear effectiveness and ready detection of the mobile dsRNA pool (Fig. S5), primary siRNAs (diagnosed by the *GUS* portion of the dsRNA; Fig. S5) were absent for TMV and PVX infected plants. This was despite the presence of readily detectable *GFP* siRNAs (at least in the case of TMV; Fig. S5), indicating that initial siRNA production is likely occurring in a very precise spatio-temporal manner.

Here we demonstrate for the first time that topically applied dsRNA can move and act non-cell autonomously in multiple plant species and within a variety of key tissue types and against both plant pathogenic fungi and viruses. Importantly, we demonstrated that dsRNA moves as an intact unprocessed molecule which can be maintained over time in newly developed tissue. These facets of movement have several important benefits for pathogen management. Topically applied dsRNA is known to be rapidly degraded when added to soil, making the control of root-based pathogens difficult. Our finding that dsRNA has the ability to move from leaves to roots could stand to circumvent this problem. Additionally, the finding that dsRNA moves within the apoplast rather than the symplast has several implications for pathogen management initiated by this form of RNAi-based movement. Viruses are known to not only move via symplastic channels in plants, but also to manipulate and alter plasmodesmata in the process (26). Callose deposition is also used as a defence mechanism during fungal infection (27, 28). By travelling via the apoplast, dsRNA can move independently of any virus movement or virus/fungal mediated plasmodesmata manipulation. As a result, dsRNA has the potential of forming an independent resistance front. Also, by remaining in the apoplast and not entering the cell prior to infection, the plant can maintain a ‘pool’ of dsRNA primed for processing upon pathogen challenge. This is backed up by the lack of evidence for siRNA production either in the application (5) or destination tissue ((10); this work) in either unchallenged or challenged plants. The difficulty of detecting siRNAs indicates a highly precise spatio-temporal production of siRNAs. Indeed, we were unable to detect primary siRNAs in any of our topical application, virus movement or fungal infection experiments, despite of their clear functionality and the ready detection graft transmissible endogenous siRNA movement (Fig. S5 and S6). These key molecular events will have to be deciphered in future if the use of topically applied mobile dsRNA is to reach its true potential in a wide range of important crop species. However, given that we see effective targeting of experimental viral and fungal systems, which themselves are several orders of magnitude in excess of anything encountered in nature, the use of this mobile immune system has enormous potential for crop protection in the coming years.

## Materials and Methods

### Plant materials

Unless otherwise stated all wild-type and mutant plants are in the Columbia (Col-0) background. *The dcl2-1 dcl3-1 dcl4-2* (29) mutant, *pSUC::SUL* (SS) (22) and GFP RNAi (30) transgenic lines have been described previously. Canola (*Brassica napus*) seeds were a gift from Professor Ian Goodwin (The University of Queensland). Tomato plants were the microtom variety and *Nicotiana benthamiana* plants were the common lab variety.

### Plant growth conditions

Unless specifically stated, all plants were germinated and grown under a 12h:12h light:dark cycle at 22°C on soil or 0.5x Murashige and Skoog (MS) media. For experiments relating to roots, plants were first germinated on 0.5x MS media and grown vertically for 1-2 weeks. Seedlings were transferred to end-cut 0.5 mL tubes filled with 1% plant agar contained within a 1 mL tip box for hydroponic growth. Plants were supplemented with 500 mL of 0.5x Cramer’s solution (0.75 mM Ca(NO_3_)_2_, 0.625 mM KNO_3_, 0.5 mM (NH_4_)_2_SO_4_, 375 μM MgSO_4_,, 50 μM Na_2_O_3_Si, 36 μM Na[Fe(EDTA)], 25 μM KCl, 25 μM H_3_BO_3_, 5 μM MnSO_4_, 1 μM ZnSO_4_, 0.75 μM CuSO_4_ and 0.0375 μM (NH_4_)_6_Mo_7_O_24_). Nutrient solution was exchanged every 1-2 weeks. For phosphate starvation (-Pi) the same 0.5x Cramer’s media was used but the 250 μM KH_2_PO_4_ was omitted.

### dsRNA production

Typically, dsRNA was produced using the HiScribe T7 High yield RNA synthesis kit (New England BioLabs). Templates were PCR products with T7 promoter sequences added to both ends to facilitate bidirectional transcription using the oligonucleotides listed in Table S1. Products were amplified from reverse transcribed RNA from the desired organism. For most experiments dsRNA was synthesised using a ~350 nt sequence of *GFP* using the primers GFP-T7-F and GFP-T7-R. For experiments involving PVX-GFP and TMV-GFP targeting, overlapping fusion PCRs were used to generate dsRNA PCR templates containing ~350 nt of *GFP* sequence at the 5’ and ~250 nt of *GUS* at the 3’ end. Primers used to generate either *mGFP-GUS* (PVX-GFP targeting) or *eGFP-GUS* (TMV-GFP targeting) are listed in Table S1. All *in vitro* synthesised dsRNA was treated with turbo DNase (Invitrogen), purified using TRIsure reagent (Bioline Meridian Biosystems), denatured and annealed prior to use.

### Topical application of dsRNA

For all experiments involving molecular analysis of movement, *in vitro* synthesis dsRNA was diluted to a concentration of 300 ng/μL and supplemented with penetrant at a concentration of 0.01%. For Arabidopsis, 3-5 μL was added per leaf and for canola, tomato and *N.benthamiana*, 10-20 μL was added per leaf.

### Petiole application of dsRNA

Petiole absorption of dsRNA was performed as described previously (10). RNA was adjusted to a concentration of 250 ng/μL and 200 μL was added to the lid of a closed 0.5 μL tube which had the base cut off. The 2^nd^ or 3^rd^ leaf of ~3-week-old *N. benthamiana* plants was removed to as close to the end of the petiole as possible. The exposed petiole was then inserted into the tube opening and sealed with parafilm.

### Micrografting

Micrografting was performed as described previously (30, 31). Briefly, seeds were germinated on vertical plates containing 0.5x MS media and grown vertically for 5-6 days. Grafts were performed by cutting with a no.15 scalpel blade across the hypocotyl ~ 1 mm from the shoot apex and joining with a rootstock cut in an identical fashion. Grafted seedlings were aligned under a dissecting microscope on a single 0.45 μM nitrocellulose filter (Millipore) placed on top of two layers of moistened Whatman filter paper in a 90 mm Petri dish. Grafts were maintained in a vertical orientation for 7 days under long day conditions. Successful grafts were subsequently transferred to the hydroponic growth setup described above.

### Meselect isolation of vascular tissue

Separation of epidermal and vascular tissue (Meslect) of rosette leaves was performed as described previously (12). Leaves were secured between two pieces of tape with abaxial epidermal cells physically separated from the remaining cell types (including vascular tissue) by peeling the tape apart. Both the upper (containing the vasculature) and the lower (abaxial epidermal cells) epidermis were incubated in protoplasting solution (1% cellulase Onozuka RS, 0.4 M mannitol, 10 mM CaCl_2_, 20 mM KCl, 0.1% BSA and 20 mM MES, pH 5.7) with gentle agitation. The lower epidermal side and the upper cells/vasculature were incubated for 15 min and 20 min respectively. Post-incubation in protoplasting solution, both tissue types were washed twice with washing solution (154 mM NaCl, 125 mM CaCl_2_, 5 mM KCl, 5 mM glucose and 2 mM MES (pH 5.7)). After washing, the epidermal tape was frozen in liquid nitrogen and the vasculature was removed from the tape and also snap frozen. Tissue was stored at −80°C until RNA extraction as described below.

### Apoplast and extracellular vesicle extraction

Apoplastic fluids were purified from leaves of tomato or *N. benthamiana* or roots of Arabidopsis plants by first syringe infiltrating tissue with infiltration buffer (20 mM MES, 2 mM CaCl_2_, 0.1 M NaCl, pH 6.0). Apoplastic fluid was subsequently extracted via low-speed centrifugation (900*g*, 4°C). Extracts were then spun at 2,000*g* for 30 min, filtered through 0.45 μm columns and then spun at 10,000*g* for a further 30 min. The resulting supernatant was used as apoplastic fluid. Extracellular vesicles were purified from apoplastic fluid by spinning at 100,000*g* for 1 h (24). The resulting pellet was washed with infiltration buffer and spun for another hour at 100,000*g* prior to extraction of RNA.

### Fungal material and infection

*Fusarium oxysporum* (Fo5176) transformed with a *pGPDA-GFP* binary vector as previously described (32). *Verticillium dahliae* expressing *pTEF-GFP* has been described previously (33). Spores were isolated either from plates (1/2 Potato Dextrose Agar) or liquid cultures grown in Potato Dextrose Broth. Spore density was calculated using a haemocytometer and adjusted to a spore count of between 1 x 10^6^ and 3 x 10^6^ spores/mL prior to infection. The roots of hydroponically grown plants were exposed to spores at the indicated density for 30-60 s. Plants were then grown on 1% agar for 24 h prior to being transferred back to hydroponic conditions until harvesting.

### Virus material and infection

Virus material and infections with TMV-GFP were performed essentially as described previously (34) using *Agrobacterium tumefaciens*-based leaf infiltration. Briefly, *Agrobacterium* cultures were grown to an OD_600_ of 0.5, pelleted and resuspended in a buffer containing 10 mM MgCl_2_, 10 mM MES (pH 5.6) and 100 μM acetosyringone followed by an incubation at room temperature for a minimum of 2 h. Infections were performed 72 h post treatment with dsRNA. Assessment of infections via GFP imaging and molecular analysis were performed 2-3 days post infiltration. The *PVX-GFP* binary vector was described previously (35). Virus inoculum consisted of systemically infected PVX-GFP *N. benthamiana* leaves stored at −80°C. Leaf material was homogenized in 0.1 M NaPO_4_ buffer (pH 7) containing 1% celite and rub inoculations were carried out 3-4 days post dsRNA treatment. Infections were assessed via phenotypic and molecular analysis 4 days post infection.

### GFP imaging and quantification

GFP imaging of virus-infected plants was performed essentially as described previously (36). Camera settings were altered depending on the experiment but remained unchanged within a single one. Quantification of GFP fluorescence was performed by collecting a small disc of tissue (8 mm diameter) using a cork borer, snap freezing in liquid nitrogen and disrupting tissue using a TissueLyser (Eppendorf). Tissue was subsequently resuspended in 150 μL of lysis buffer (50 mM Tris-HCl pH 7.5, 150 mM NaCl, 1 mM EDTA and 1% NP-40). After a brief spin (1,000*g* for 1 min), 100 μL was transferred to fresh tubes. Fluorescence was quantified using a Glomax Discover microplate reader (Promega) using an excitation wavelength of 475 nm and emission was captured at 500-550 nm.

### RNA gel blot analysis

Total RNA was extracted by grinding snap frozen tissue to a fine powder in liquid nitrogen with a motor and pestle using TRIsure reagent (Bioline Meridian Biosystems), following the manufacturer’s instructions. For high molecular weight RNA gel blot analysis, total RNA (1 – 10 μg) was separated using formaldehyde (2.2 M) gel electrophoresis and transferred to a Hybond-NX Nylon membrane (GE Healthcare) via capillary transfer. After transfer, RNA was immobilized using UV crosslinking (125 mJ/cm^2^) and hybridized at 42°C using Abion ULTRAhyb hybridization buffer and PCR DIG probes (Roche) synthesised according to the manufacturer’s instructions. Membranes were washed twice with low stringency wash buffer (2x Saline-Sodium Citrate (SSC), 0.1% SDS) for 10 min followed by one wash each in high stringency (0.5x SSC, 0.1% SDS) and ultra-high stringency buffer (0.1x SSC, 0.1% SDS) each for 10 min. Membranes were then blocked using 1% blocking solution (Roche) in 1X maleic acid buffer (pH 7.5) for 30 min, and probes detected using a 1:10,000 dilution of Anti-DIG antibody (Roche) for a further 60 min in 1% blocking buffer. Membranes were washed in 1 X maleic acid buffer containing 0.3% (v/v) Tween 20, three times each for 15 min. Membranes were then equilibrated for 5 min in detection buffer (0.1 M Tris pH 9.5, 0.1M NaCl and detected using CDP star-ready to use (Sigma) and revealed using either X-ray film or an iBright detection system (Thermo Fisher). Membranes were stripped using stripping solution (50 mM Tris pH 7.4, 50% formamide, 5% SDS) for 1 h at 60°C prior to re-probing.

Low molecular weight northern blots were performed essentially as described previously (13). Total RNA (5 – 20 μg) was separated on a 17% denaturing polyacrylamide gel containing 8 M urea. RNA was electro-transferred to a Hybond-NX nylon membrane (GE Healthcare) and immobilized by N-Ethyl-N’-(dimethylaminopropyl) carbodiimide (EDC)- based cross-linking as previously described (37). Probes were generated using either oligonucleotides complementary to known siRNA or miRNA sequences or combined longer oligonucleotide probes covering ~ 150 nt of dsRNA sequences giving rise to siRNA. Probes were labelled with the DIG Oligonucleotide 3’-End Labelling Kit 2^nd^ generation (Roche) with probe sequences listed in Table S1. Hybridizations were carried out at 42 °C using Ambion ULTRAhyb-oligo buffer for a minimum of 2 h. Membranes were washed 3x 10 min in 4x SSC and antibody detection performed as described above.

### Real-time qRT-PCR analysis

Total RNA extracted from was extracted using TRIsure reagent (Bioline Meridian Biosystems). Total RNA (1-5 μg) was treated with Turbo DNase (Invitrogen) and reverse-transcribed using Tetro Reverse Transcriptase (Bioline Meridian Biosystems) following the manufacturer’s instructions. Real-time quantitative reverse transcriptase PCR (RT-qPCR) was performed using a Biorad CFX384 Optics Module with SensiFAST SYBER No-ROX (Bioline Meridian Biosystems) using gene-specific primers listed in Supplementary Table 1. Fusarium GFP transcripts were normalised to glyceraldehyde 3-phosphate dehydrogenase controls and Verticillium GFP transcripts were normalised to tubulin. RT-qPCR was carried out using technical triplicates using the following cycling conditions: 95 °C for 3 min, followed by 40 cycles of 95 °C for 10 s, 60°C for 10 s and 72°C for 20 s. A melt curve was performed at the end of every run to confirm oligo specificity. Threshold cycle values (C_t_) were determined by calculating the second-derivative maximum of three technical triplicates for each biological sample. Data were analysed using Prism-GraphPad Software v8.4.0.

## Acknowledgments

We thank Mitter lab members for fruitful discussions and Dr Lisa Wittenhagen for valuable feedback on the manuscript. Dr Donald Gardiner for the GFP expressing Verticillium and Fusarium strains and Prof Olivier Voinnet for the *SS* seeds. New England Biolabs Australia are thanked for molecular biology reagents. This research was supported by the Australian Research Council Research Hub for Sustainable Crop Protection (project number IH190100022) and the Australian Government.

## Author Contributions

C.A.B designed the project and all experiments with input from A.S. and F.F.F. and N.M.. C.A.B., A.S. and F.F.F. performed all experiments. P.M.W. and B.J.C contributed reagents and experimental space. C.A.B., A.S. and F.F.F. interpreted results. C.A.B assembled all of the figures and wrote the manuscript. All authors read and approved the manuscript.

## References

1. S.-W. Ding, O. Voinnet, Antiviral immunity directed by small RNAs. Cell 130, 413–426 (2007).

2. K. H. J. Gordon, P. M. Waterhouse, RNAi for insect-proof plants. Nat Biotechnol 25, 1231–1232 (2007).

3. C. A. Brosnan, O. Voinnet, Cell-to-cell and long-distance siRNA movement in plants: mechanisms and biological implications. Curr. Opin. Plant Biol. 14, 580–587 (2011).

4. F. Tenllado, J. R. Díaz-Ruíz, Double-stranded RNA-mediated interference with plant virus infection. J Virol 75, 12288–12297 (2001).

5. N. Mitter, et al., Clay nanosheets for topical delivery of RNAi for sustained protection against plant viruses. Nat Plants 3, 16207 (2017).

6. S. J. Fletcher, P. T. Reeves, B. T. Hoang, N. Mitter, A Perspective on RNAi-Based Biopesticides. Frontiers in Plant Science 11, 51 (2020).

7. M. Wang, et al., Bidirectional cross-kingdom RNAi and fungal uptake of external RNAs confer plant protection. Nat Plants 2, 16151 (2016).

8. O. Christiaens, S. Whyard, A. M. Vélez, G. Smagghe, Double-Stranded RNA Technology to Control Insect Pests: Current Status and Challenges. Front. Plant Sci. 11, 451 (2020).

9. A. Koch, et al., An RNAi-Based Control of Fusarium graminearum Infections Through Spraying of Long dsRNAs Involves a Plant Passage and Is Controlled by the Fungal Silencing Machinery. PLoS Pathog 12, e1005901 (2016).

10. A. Dalakouras, et al., Delivery of Hairpin RNAs and Small RNAs Into Woody and Herbaceous Plants by Trunk Injection and Petiole Absorption. Front Plant Sci 9, 1253 (2018).

11. D. Biedenkopf, et al., Systemic spreading of exogenous applied RNA biopesticides in the crop plant Hordeum vulgare. ExRNA 2, 12 (2020).

12. J. Svozil, W. Gruissem, K. Baerenfaller, Meselect - A Rapid and Effective Method for the Separation of the Main Leaf Tissue Types. Front Plant Sci 7, 1701 (2016).

13. C. A. Brosnan, et al., Genome-scale, single-cell-type resolution of microRNA activities within a whole plant organ. EMBO J. 38, e100754 (2019).

14. E. A. Devers, et al., Movement and differential consumption of short interfering RNA duplexes underlie mobile RNA interference. Nat. Plants (2020) https://doi.org/10.1038/s41477-020-0687-2 (July 7, 2020).

15. A. Maizel, K. Markmann, M. Timmermans, A. Wachter, To move or not to move: roles and specificity of plant RNA mobility. Current Opinion in Plant Biology 57, 52–60 (2020).

16. O. Voinnet, P. Vain, S. Angell, D. C. Baulcombe, Systemic spread of sequence-specific transgene RNA degradation in plants is initiated by localized introduction of ectopic promoterless DNA. Cell 95, 177–187 (1998).

17. K. Kobayashi, P. Zambryski, RNA silencing and its cell-to-cell spread during Arabidopsis embryogenesis. Plant J. 50, 597–604 (2007).

18. T. Rosas-Diaz, et al., A virus-targeted plant receptor-like kinase promotes cell-to-cell spread of RNAi. Proc. Natl. Acad. Sci. U.S.A. 115, 1388–1393 (2018).

19. A. Carlsbecker, et al., Cell signalling by microRNA165/6 directs gene dose-dependent root cell fate. Nature 465, 316–321 (2010).

20. A. Vatén, et al., Callose biosynthesis regulates symplastic trafficking during root development. Dev. Cell 21, 1144–1155 (2011).

21. J. Müller, et al., Iron-Dependent Callose Deposition Adjusts Root Meristem Maintenance to Phosphate Availability. Developmental Cell 33, 216–230 (2015).

22. C. Himber, P. Dunoyer, G. Moissiard, C. Ritzenthaler, O. Voinnet, Transitivity-dependent and -independent cell-to-cell movement of RNA silencing. EMBO J. 22, 4523–4533 (2003).

23. P. Baldrich, et al., Plant Extracellular Vesicles Contain Diverse Small RNA Species and Are Enriched in 10-to 17-Nucleotide “Tiny” RNAs. Plant Cell 31, 315–324 (2019).

24. Q. Cai, et al., Plants send small RNAs in extracellular vesicles to fungal pathogen to silence virulence genes. Science 360, 1126–1129 (2018).

25. K. Necira, et al., Topical Application of Escherichia coli-Encapsulated dsRNA Induces Resistance in Nicotiana benthamiana to Potato Viruses and Involves RDR6 and Combined Activities of DCL2 and DCL4. Plants (Basel) 10, 644 (2021).

26. M. Heinlein, Plasmodesmata: channels for viruses on the move. Methods Mol Biol 1217, 25–52 (2015).

27. E. Luna, et al., Callose deposition: a multifaceted plant defense response. Mol Plant Microbe Interact 24, 183–193 (2011).

28. M.-C. Caillaud, et al., The plasmodesmal protein PDLP1 localises to haustoria-associated membranes during downy mildew infection and regulates callose deposition. PLoS Pathog 10, e1004496 (2014).

29. A. Deleris, et al., Hierarchical action and inhibition of plant Dicer-like proteins in antiviral defense. Science 313, 68–71 (2006).

30. C. A. Brosnan, et al., Nuclear gene silencing directs reception of long-distance mRNA silencing in Arabidopsis. Proc. Natl. Acad. Sci. U.S.A. 104, 14741–14746 (2007).

31. C. G. N. Turnbull, J. P. Booker, H. M. O. Leyser, Micrografting techniques for testing long-distance signalling in Arabidopsis. Plant J. 32, 255–262 (2002).

32. B. N. Kidd, et al., Auxin Signaling and Transport Promote Susceptibility to the Root-Infecting Fungal Pathogen *Fusarium oxysporum* in *Arabidopsis*. MPMI 24, 733–748 (2011).

33. R. Sabburg, et al., “A method for high throughput image based antifungal screening” (Microbiology, 2021) https://doi.org/10.1101/2021.08.18.456906 (October 5, 2021).

34. J. Bally, et al., The extremophile Nicotiana benthamiana has traded viral defence for early vigour. Nat Plants 1, 15165 (2015).

35. A. F. Fusaro, et al., The Enamovirus P0 protein is a silencing suppressor which inhibits local and systemic RNA silencing through AGO1 degradation. Virology 426, 178–187 (2012).

36. F. F de Felippes, et al., The key role of terminators on the expression and post-transcriptional gene silencing of transgenes. Plant J 104, 96–112 (2020).

37. G. S. Pall, A. J. Hamilton, Improved northern blot method for enhanced detection of small RNA. Nat Protoc 3, 1077–1084 (2008).

