## Supplementary Information for "Intact double stranded RNA is mobile and triggers RNAi against viral and fungal plant pathogens"

**Supplementary Figure legends**

**Supplementary Figure 1. Arabidopsis displays low levels of dsRNA movement to young leaf tissue.** Northern blot analysis of dsRNA movement from application leaf to young leaves 24 h post application. Note that this is the same membrane used in Figure 1a. Ribosomal RNA (rRNA) serves as a loading control.

**Supplementary Figure 2. Limited movement of dsRNA in *Nictotiana benthamiana*. a,** Northern blot detection of dsRNA in applied leaf and sequential leaves distal from application point (leaf 1-3). Two exposures are shown for the *GFP* probe. Ribosomal RNA (rRNA) serves as a loading control.

**Supplementary Figure 3. Blocking of symplastic siRNA movement via constitutive callose deposition.** **a**, Representative photographs of ~5-week-old SS control and phosphate starved leaves. Leaves were photographed two weeks post-treatment and represent ones which emerged post-treatment. Scale bars represent 1 cm **b**, Quantitative RT-PCR of phosphate responsive marker SQD-1 normalized to actin2 in control or -Pi treated plants. Error bars represent +/- s.e.m. of 3 biological replicates. *P* value represents a two-way unpaired students *t*-test. **c**, RNA gel blot analysis of *SS* (SUL) siRNA levels in post-treatment lines. U6 serves as a loading control.

**Supplementary Figure 4. Lack of siRNAs in distal mobile dsRNA receiving root tissue in Arabidopsis.** Small RNA gel blot analysis of GFP RNAi, GFP RNAi grafted as a scion onto *dcl234* rootstocks, WT and *dcl234* roots where *GFP* dsRNA was applied to the leaves and non-dsRNA treated WT and *dcl234* plants. Two GFP exposures are shown along with graft transmissible endo-siRNA (rep2). miR159 serves as a non-mobile miRNA and U6 as loading controls respectively.

**Supplementary Figure 5. Full length dsRNA and siRNA analysis in virus infected *N. benthamiana* plants. a**, RNA gel blot analysis of dsRNA from leaves used in TMV-GFP experiments depicted in Figure 4f. **b**, Small RNA gel blot analysis of RNA from the mobile leaf used in TMV-GFP experiments depicted in Figure 4f. 159 and U6 serve as endogenous miRNA and loading controls respectively **c**, RNA gel blot analysis of dsRNA from mobile leaves used in PVX-GFP experiments depicted in Figure 4g. **b**, Small RNA gel blot analysis of RNA from the mobile leaf used in PVX-GFP experiments depicted in Figure 4g. * depicts non-specific band present in all samples. 159 and U6 serve as endogenous miRNA and loading controls respectively.

**Supplementary Figure 6. Assaying siRNAs in Arabidopsis plants infected with fungal pathogens. a,** Small RNA gel blot analysis of GF RNAi, GF RNAi grafted as scion onto *dcl234* rootstock, WT roots (infected and mock) where dsRNA was applied to the leaves and non-treated WT infected plants. Fungal infections were with *Fusarium oxysporum* (f.ox). Two GFP exposures are shown along with graft transmissible endo-siRNA (rep2). miR159 serves as a non-mobile miRNA and U6 as loading controls respectively. **b,** Small RNA gel blot analysis of GF RNAi, GF RNAi grafted as scion onto *dcl234* rootstock, WT roots (infected and mock) where dsRNA was applied to the leaves and non-dsRNA treated WT infected plants. Fungal infections were with *Verticillium dahliae* (v.d). Two GFP exposures are shown along with graft transmissible endo-siRNA (rep2). miR159 serves as a non-mobile miRNA and U6 as loading controls respectively.

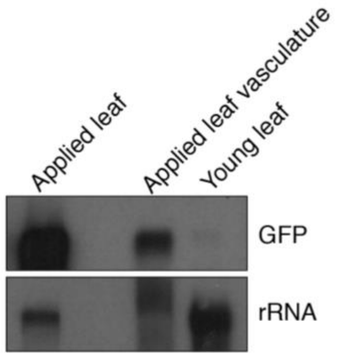


**Supplementary Figure 1**

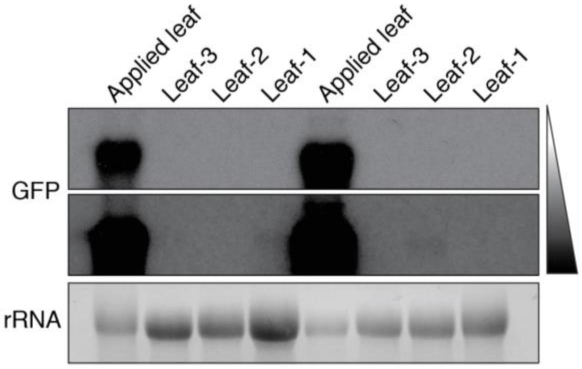


**Supplementary Figure 2**

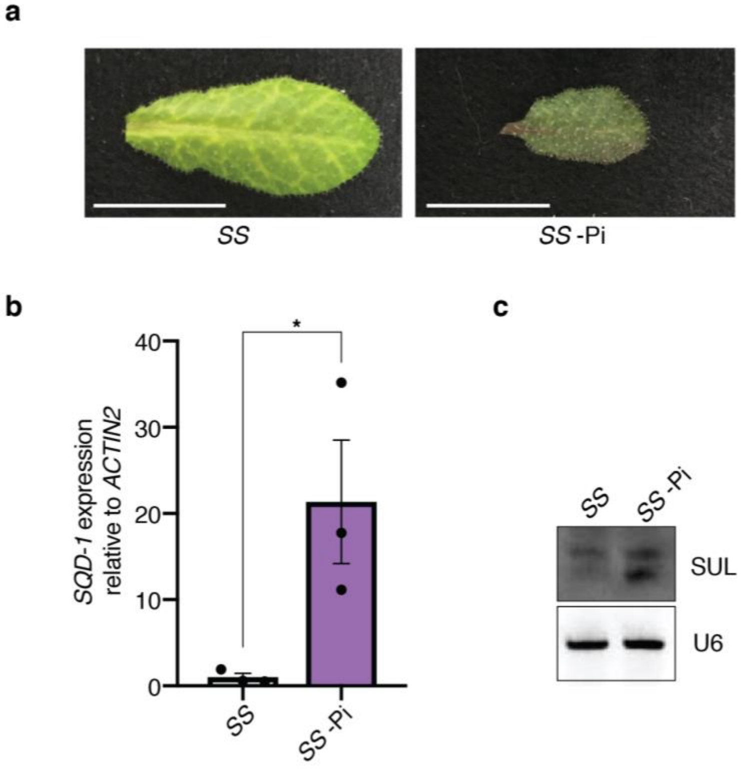


**Supplementary Figure 3**

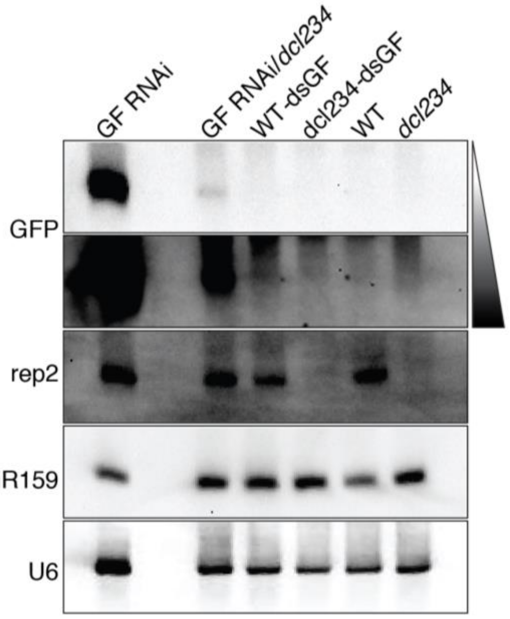


**Supplementary Figure 4**

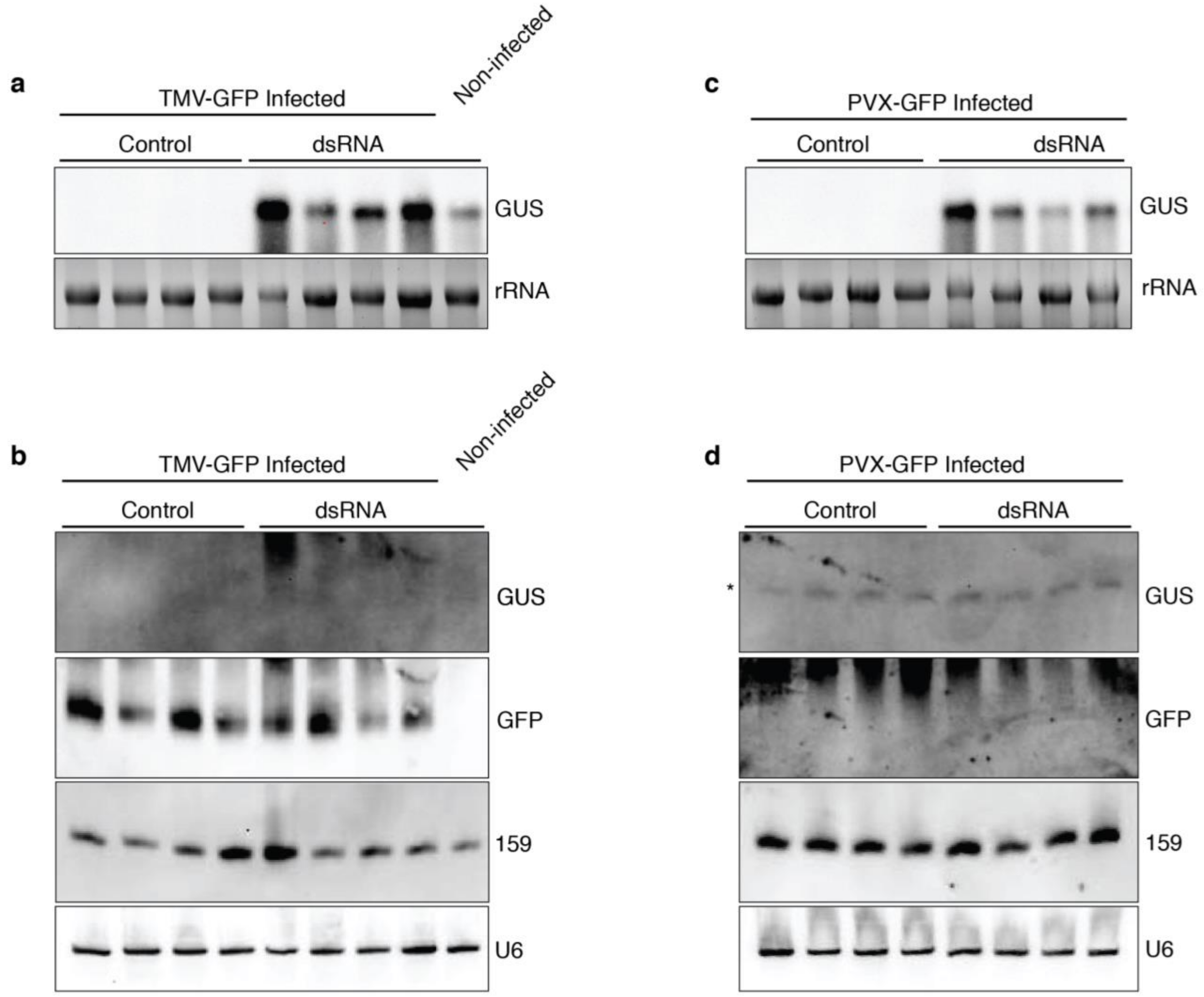


**Supplementary Figure 5**

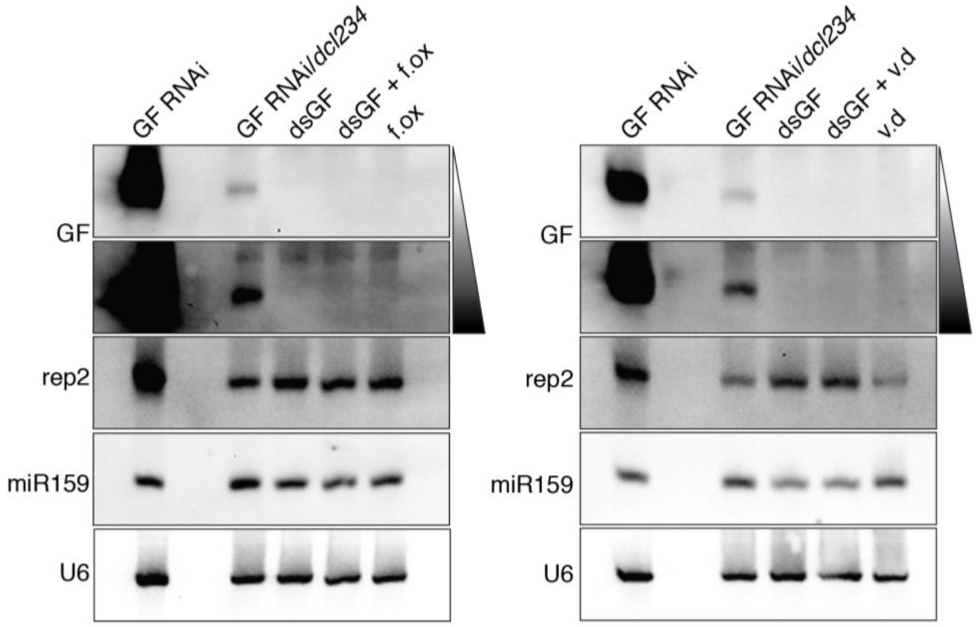


**Supplementary Figure 6**

**Supplementary Table 1:** List of primers used for this study

| | Name | Sequence (5' to 3') |
|---|---|---|
| **dsRNA template** | **GFP-T7-F** | TAATACGACTCACTATAGGGAGGACGACGGCAACTACAAG |
| | **GFP-T7-R** | TAATACGACTCACTATAGGGTCTCGTTGGGGTCTTTGCTC |
| | **ENO-T7-F** | TAATACGACTCACTATAGGGCCACGGGCGAGTTCGAGGCC |
| | **ENO-T7-R** | TAATACGACTCACTATAGGGATAATGAACTCCTGCATCGC |
| | **mG-F** | ATGAGTAAAGGAGAAGAACT |
| | **mG-GUS-R** | GGTTTCTACAGGACGTAACATGAAAAATATAGTTCTTTCCTGTAC |
| | **GUS-F** | ATGTTACGTCCTGTAGAAACC |
| | **GUS-R** | ACGTTGCCCGCATAATTACGA |
| | **mG-T7-F** | TAATACGACTCACTATAGGGATGAGTAAAGGAGAAGAACT |
| | **GUS-T7-R** | TAATACGACTCACTATAGGGACGTTGCCCGCATAATTACGA |
| | **eG-F** | ATGGTGAGCAAGGGCGAGGAG |
| | **eG-GUS-R** | GGTTTCTACAGGACGTAACATGAAGATGGTGCGCTCCTGGA |
| | **eG-T7-F** | TAATACGACTCACTATAGGGATGGTGAGCAAGGGCGAGGAG |
| **Probes** | **GF-1** | GTGCCCATCCTGGTCGAGCTGGACGGCGACGTAAACGGCCACAAGTTCAGCGTGTCCGGC |
| | **GF-2** | GTAAACGGCCACAAGTTCAGCGTGTCCGGCGAGGGCGAGGGCGATGCCACCTACGGCAAG |
| | **GF-3** | GAGGGCGAGGGCGATGCCACCTACGGCAAGCTGACCCTGAAGTTCATCTGCACCACCGGC |
| | **GF-4** | CTGACCCTGAAGTTCATCTGCACCACCGGCAAGCTGCCCGTGCCCTGGCCCACCCTCGTG |
| | **SUL-1** | CTTTGAGCCTGGTTTGTTGGCTAAAGCTAATAGAGGGATTCTTTATGTTGATGAAGTTAA |
| | **SUL-2** | TAGAGGGATTCTTTATGTTGATGAAGTTAATCTCTTGGATGATCATTTGGTTGATGTTCT |
| | **SUL-3** | TCTCTTGGATGATCATTTGGTTGATGTTCTTTTGGATTCAGCTGCTTCTGGTTGGAATAC |
| | **SUL-4** | TTTGGATTCAGCTGCTTCTGGTTGGAATACGGTTGAGAGAGAAGGGATTTCGATTTCTCA |
| | **GUS-1** | ATGTTACGTCCTGTAGAAACCCCAACCCGTGAAATCAAAAACTCGACGGCCTGTGGGCA |
| | **GUS-2** | GAATCAAAAACTCGACGGCCTGTGGGCATTCAGTCTGGATCGCGAAAACTGTGGAATT |
| | **GUS-3** | TTCAGTCTGGATCGCGAAAACTGTGGAATTGATCAGCGTTGGTGGGAAAGCGCGTTACAA |
| | **GUS-4** | GATCAGCGTTGGTGGGAAAGCGCGTTACAAGAAAGCCGGGCAATGCTGTGCCAGGCAGT |
| | **rep2** | GCGGGACGGGTTTGGCAGGACGTTACTTAAT |
| | **miR159** | TAGAGCTCCCTTCAATCCAAA |
| | **U6** | GTTTTATCAAGTCCCAGACCGTATCAAATAT |
| | **25S** | CCTCCGCTTATTGATATGCTTAAACTCAGCGGGTAATCCCGCCTGA |
| **q-RT-PCR** | **GFP-RT-3-F** | TGAGCAAAGACCCCAACGAG |
| | **GFP-RT-3-R** | GTCCATGCCGTGAGTGATCC |
| | **f.ox-GPD-RT-F** | AAGGGTGCTTCTTACGACC |
| | **f.ox-GPD-RT-R** | ATCGGAGGAGACAACATCG |
| | **SQD-1-RT-F** | CAGCAAGAATTCAGTTAAGCC |
| | **SQD-1-RT-R** | GTTTCTTCTCTTGGAAGAGAAG |
| | **v.d-tub-RT-F** | TCCACCTTCGTCGGTAACTC |
| | **v.d-tub-RT-R** | GCCTCCTCCTCGTACTCCTC |
| | **act-2-F** | GGCACCCTGTTCTTCTTACCG |
| | **act-2-R** | AACCCTCGTAGATTGGCACA |
